# A chromosome-level genome of the giant vinegaroon *Mastigoproctus giganteus* exhibits the signature of pre-Silurian whole genome duplication

**DOI:** 10.1101/2024.11.08.622695

**Authors:** Siddharth S. Kulkarni, Benjamin C. Klementz, Prashant P. Sharma

## Abstract

Within the arachnids, chromosome-level genome assemblies have greatly accelerated the understanding of gene family evolution and developmental genomics in key groups, such as spiders (Araneae), mites and ticks (Acariformes and Parasitiformes). Among other poorly studied arachnid orders that lack genome assemblies altogether are the clade Pedipalpi, which is comprised of three orders that form the sister group of spiders, which diverged over 400 Mya. We close this gap by generating the first chromosome-level assembly from a single specimen of the vinegaroon *Mastigoproctus giganteus* (Uropygi). We show that this highly complete genome retains plesiomorphic conditions for many gene families that have undergone lineage-specific derivations within the more diverse spiders. Consistent with the phylogenetic position of Uropygi, macrosynteny in the *M. giganteus* genome substantiates the signature of an ancient whole genome duplication.

## 1 Introduction

Chelicerata, the sister group of the remaining arthropods, constitutes a phylogenetically significant and ancient lineage that diversified in the Cambrian (Dunlop, 2010; Giribet, 2018; Ballesteros et al., 2021). Over 120,000 described species of chelicerates are presently divided into 14 orders which vary markedly in their species richness. Whereas diverse groups such as spiders and mites bear tens of thousands of described (and large proportions of undescribed) taxa, relictual lineages such as horseshoe crabs are limited to just four living species (Pepato et al., 2010; Obst et al., 2012; Sharma, 2023b). The basis for these patterns of asymmetrical species richness has been investigated from several perspectives, such as evolutionary novelties (venoms; silks; webs), clade age, biogeography, and most recently, genome architecture. Chelicerata is notable for the incidence of multiple whole genome duplications, with three of these occurring on the branch subtending modern horseshoe crabs, and a separate pre-Silurian duplication uniting a subset of six orders called Arachnopulmonata (Kenny et al., 2016; Schwager et al., 2017; Shingate et al., 2020a; Nong et al., 2021; Ontano et al., 2021; Ballesteros et al., 2022) (Fig. 1A). Included among the arachnopulmonates are three of the four venomous arachnid orders (spiders, scorpions, and pseudoscorpions) and two of the three silk-producing orders (spiders and pseudoscorpions) (Santibáñez-López et al., 2018, 2022; Ontano et al., 2021). Some arachnopulmonate species are also renowned for independent origins of visual acuity, as exemplified by jumping spiders (Salticidae) and huntsmen spiders (Sparassidae) (Nørgaard et al., 2008; Jakob et al., 2018; Chong et al., 2024; Winsor et al., 2024).

**Figure 1.**
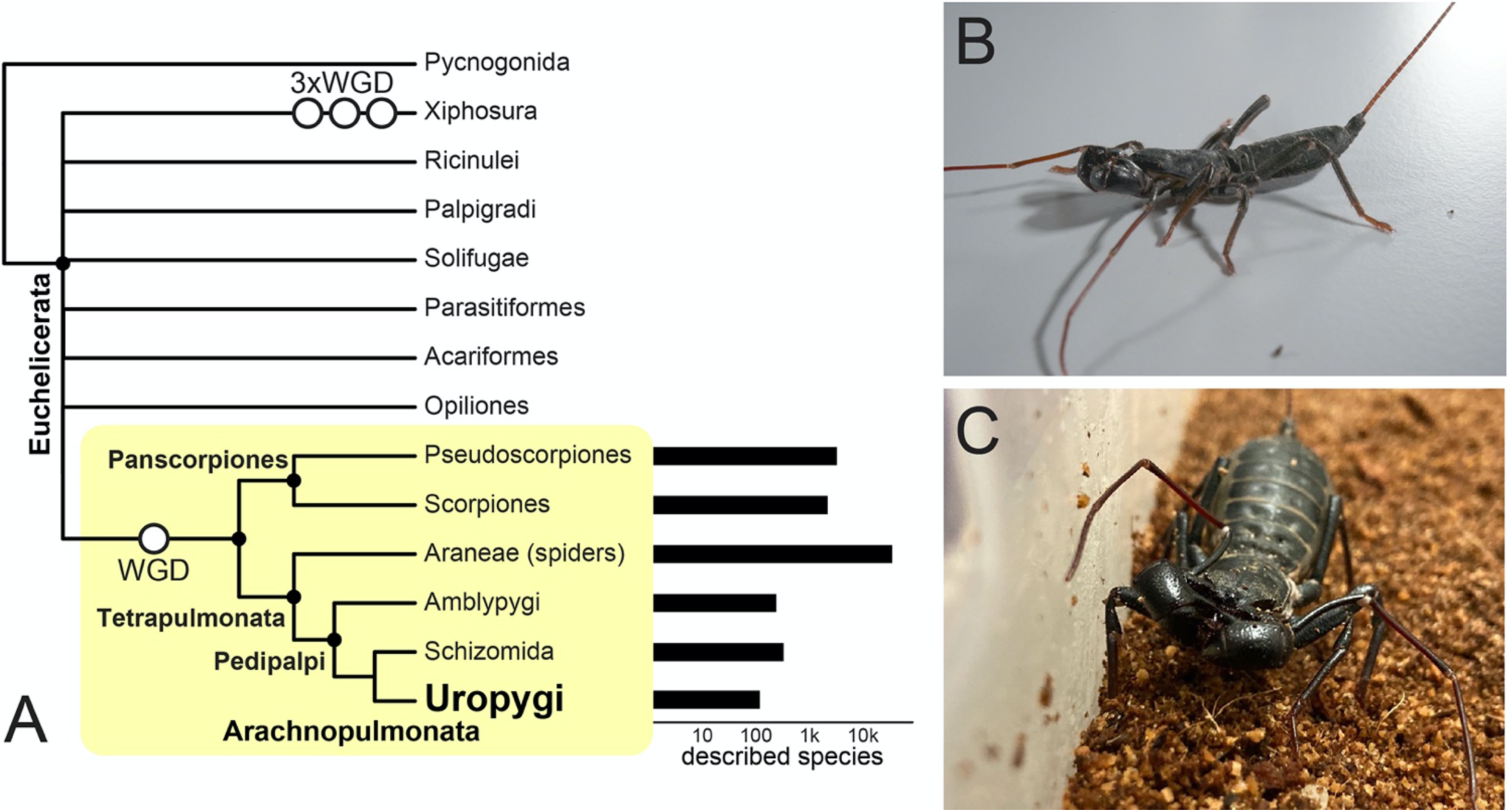
(A) Phylogeny of Chelicerata with emphasis on the relationships between Arachnopulmonata. Tree topology is based on Ballesteros et al. (2022). (B) Live habitus of *M. giganteus* in lateral view. (C) Live habitus of *M. giganteus* in frontal view.

The relationship between the phylogenetic concentration of these traits in arachnopulmonates and their whole genome duplication is not well understood. Whole genome duplication events are thought to accelerate the evolution of adaptive traits in successful groups, as new gene copies acquire new functions over time. This framework is frequently applied to explain the evolutionary success of spiders in particular (Schomburg et al., 2015; Schwager et al., 2015, 2017). However, there are two caveats to the putative link between whole genome duplication and spider diversification. First, functional evidence supporting this framework is limited in Chelicerata, in part due to the paucity of gene misexpression experiments in spider models that have disentangled the functions of ohnologs, and thereby demonstrated a clear link between new genes and new traits. As a result, many putative cases of gene subfunctionalization and neofunctionalization in arachnids are inferences based only on interpretations of expression patterns. Second, the attribution of adaptive value to specific gene copies requires a broader phylogenetic test of association between gene origin and trait origin. For example, many transcriptionally active gene duplicates initially thought to be spider-specific and invoked to explain spider traits were later shown to be widespread across arachnopulmonates (Samadi et al., 2015; Schomburg et al., 2015; Turetzek et al., 2017). Inversely, in other cases, gene duplicates thought to be widespread across arachnids and invoked to explain arachnid-wide traits were later shown to be restricted to arachnopulmonates alone (Turetzek et al., 2015; Klementz et al., 2024). These inferential mismatches between gene and trait origin reflect the asymmetrical availability of high-quality genome assemblies in chelicerates. Groups like horseshoe crabs, certain spider families, mites, and ticks have received substantially more investment in genomic resources than less studied orders, owing to phylogenetic significance and/or relevance to human disease and agriculture (Garb et al., 2018; Sharma, 2023b). This asymmetry of resource availability bears upon understanding the impact of the arachnopulmonate whole genome duplication because three depauperate orders within this group lack genome assemblies altogether (Amblypygi, Schizomida, and Uropygi). This trio of orders together comprises the clade Pedipalpi, the sister group to spiders (together forming Tetrapulmonata; Giribet et al., 2002; Shultz, 2007; Pepato et al., 2010; Sharma et al., 2014a; Ballesteros and Sharma, 2019) (Fig. 1a). Given the dynamism of spider genome evolution and widespread interest in their biology, the establishment of genomic resources for Pedipalpi is necessary for polarizing various evolutionary phenomena in spiders and Arachnopulmonata more broadly.

To bridge this gap, we pursued the establishment of a high-quality genome for the order Uropygi (vinegaroons), the least diverse order of Arachnopulmonata and arguably the least studied from the perspective of phylogenetics and genomics (Garb et al., 2018). Vinegaroons are typically large-bodied arachnids with two pairs of book lungs, large subchelate raptorial palps, and a posterior flagellum with numerous articles (Shultz, 1990; Schmidt, 2009). Due to their nocturnal and burrowing habits, challenges in field collection, and long generation time, comparatively little is known about vinegaroon biology (Schmidt, 2009; Schmidt et al., 2021; Schmidt and Schmidt, 2022). They are distinguished from other arachnid orders by their namesake, a pygidial gland that produces a defensive volatile spray largely composed of acetic acid (Schmidt et al., 2000). To date, 126 species have been described and the small fossil record of this group includes representatives from the Carboniferous period. These fossils largely resemble extant species, supporting the inference of long-term morphological stasis in the group (Dunlop, 2010; Knecht et al., 2024; Santana et al., 2024). Like other members of Pedipalpi, the first walking leg of vinegaroons is elongate, gracile, and endowed with numerous subdivisions of the tarsus called tarsomeres, resulting in an antenniform leg (largely used for sensory function rather than walking). All Uropygi are thought to mate after an elaborate courtship ritual lasting many hours (a derivation of the promenade a deux found across arachnopulmonates; Ontano et al., 2021; Schmidt et al., 2021). Females brood their embryos on the underside of the opisthosoma in a brood sac, and remain in a burrow with the eggs until the first instar and subsequent dispersal of the offspring. Comparatively little is known about their embryology and development (Yoshikura, 1961; Weygoldt, 1971). Notable features of embryogenesis include a downturned opisthosoma that is also observed in embryos of basally branching spider families; and inversion, an atypical process wherein the embryo splits at the ventral midline before dorsal closure (characteristic of Tetrapulmonata; Setton et al., 2019). The systematics of the group is largely based on morphological study alone (Harvey, 2002, 2007; Barrales-Alcalá et al., 2018). Molecular phylogenetic studies of vinegaroons are limited to a single study with sparse sampling of the four constituent subfamilies (Clouse et al., 2017). The phylogenetic placement of the order is well understood and highly stable, despite the present limitation of transcriptomic datasets to two species (Ballesteros et al., 2022).

Here, we generated a chromosomal-level genome assembly for a single specimen of *Mastigoproctus giganteus*, arguably the best studied species of Uropygi from the perspective of organismal biology (Weygoldt, 1971; Shultz, 1993; Schmidt et al., 2000). We demonstrate that Uropygi retain the signature of the arachnopulmonate whole genome duplication, as evidenced by systemic duplication of gene families and analyses of macrosynteny. We anticipate that this resource will greatly aid comparative genomic analysis of Chelicerata, with emphasis on polarizing genomic characters with respect to spiders.

## 2 Methods

### 2.1 PacBio HiFi library preparation and sequencing

A colony of *Mastigoproctus giganteus*, originally from a population in Hudspeth County, Texas (United States), was maintained at 28°C at the University of Wisconsin-Madison. Individuals were sexed and housed individually. Prior to DNA extraction, a single adult male was anesthetized with CO_2_ and flash frozen with liquid N_2_. Genomic DNA was extracted using Qiagen HMW DNA kit from the flash frozen *M*. *giganteus* by sectioning fragments of the appendages and extracting muscle tissue. Sequencing was performed on PacBio Sequel II, using standard manufacturer’s protocols for the Sequel II Sequencing Kit 2.0. The library was sequenced on 4 SMRT Cells (8 M) in CCS mode for 30 h. Analysis was performed with SMRT Link v10.1 software, requiring a minimum of three passes for CCS generation. Genome size and heterozygosity were estimated with a k-mer approach using Jellyfish v2.2.5 (Marçais and Kingsford, 2011) and GenomeScope (Vurture et al., 2017).

### 2.2 HiC sequencing

Omni-C library preparation and sequencing was done using the Dovetail Omni-C Kit (Cantata Bio, Scotts Valley, California) for animal tissues using the manufacturer’s protocol. Briefly, frozen tissue was thoroughly ground in a mortar with pestle in liquid N_2_, followed by fixation of chromatin with formaldehyde. The chromatin was digested with DNase I, extracted, end-repaired and ligated to a biotinylated bridge adapter followed by proximity ligation of adapter ends. Crosslinking was reversed and the DNA was purified from proteins. Biotin containing fragments were separated using streptavidin beads. The sequencing libraries tagged using unique dual indices and generated using Illumina-compatible adapters. The libraries were sequenced on an Illumina NovaSeq X+ platform, at the Biotechnology Center (University of Wisconsin, Madison) with a 2×150 bp paired end sequencing strategy. DNA and library samples were quantified using Qubit 2.0 Fluorometer (Life Technologies, Carlsbad, CA, USA) and the fragment size distribution was assessed with a TapeStation Genomic DNA ScreenTape (Agilent Technologies).

### 2.3 Genome Assembly

PacBio HiFi CCS bam reads were converted to fastq using bedtools v.2.28.0 (Quinlan and Hall, 2010) and assembled using hifiasm v.0.15.1-r329 (Cheng et al., 2021) to generate contigs. The assembly graph generated by hifiasm was converted to a set of contigs in multi-fasta format using “awk ‘/^S/{print “>“$2;print $3}’”. Additional haplotigs and contig overlaps were removed with purge_dups v1.2.5 (Guan et al., 2020). Genome length and N50 were calculated using assemblyStatistics v.1.1.3 (https://github.com/WenchaoLin/assemblyStatistics). In order to quantify biological completeness of our contig set, we used the package BUSCO v.5.4.7 (Manni et al., 2021) with the arachnid_odb10 ancestral lineage data set. We aligned the Omni-C data against the corresponding assembly with BWA-MEM (Li, 2013), found ligation junctions in Omni-C, and generated Omni-C pairs using pairtools (Open2C et al., 2024), sorting the high quality valid pairs. The resulting bam file was used to scaffold the genome using YaHS v.1.2a.1 (Zhou et al., 2023). Contact maps were generated using juicer_tools v.1.22.01 (https://github.com/aidenlab/JuicerTools). Snail plots for assembly statistics were plotted with assembly-stats v17.02.

### 2.4 Genome annotation

We extracted tissue from (1) prosomal muscles and brain and (2) the gonads of the *M. giganteus* specimen. Extractions were performed using TriZol TriReagent (Thermofisher, Waltham, Massachusetts), following manufacturer’s protocols. Libraries were prepared using the Illumina TruSeq library kit and quality control performed using a BioAnalyzer, as previously described for various arachnid species (Sharma et al., 2014a; Ballesteros and Sharma, 2019; Santibáñez-López et al., 2019). Two libraries (muscle and brain combined; gonads) were sequenced on an Illumina NovaSeq using a 2×150 bp paired end read strategy.

A library of repetitive elements was compiled using RepeatModeler2 v.2.0.5 (Flynn et al., 2020). The identified repeats were masked using RepeatMasker v.4.1.5 (Smit et al., 2015). RNASeq reads were mapped to the assemblies using HISAT2 v.2.2.1 (Kim et al., 2019). Full gene structure annotations were predicted using BRAKER3 v.3.0.3 (Gabriel et al., 2024). The gene predictions were mapped with functional annotations using eggNOG-mapper v.2 (Cantalapiedra et al., 2021) with settings to report only alignments equal or above an identity threshold of 40%, and query and subject coverage fraction thresholds of 20%. Complete protein domains were identified by searching protein sequences against the Pfam-A database (accessed 2023-09-12) using the *pfam_scan* script v.1.0. (https://github.com/aziele/pfam_scan) pipeline.

### 2.5 Identification of ohnologous gene clusters

To identify candidate genes that exhibit well-conserved clusters in bilaterian genomes, we performed BLASTP searches against the translated peptide of *M. giganteus*, using *Ixodes scapularis* (tick) and *Parasteatoda tepidariorum* (spider) sequences as queries, and retaining hits with e-values < 10^−20^. We performed searches for the constituent genes of the arthropod Hox, Nk, Iroquois, HRO, and SINE clusters, which have been well surveyed for chelicerates (Aase-Remedios et al., 2023; Sharma, 2023a). The identity of best hits was confirmed using SMART-BLAST and by gene tree analysis. For the Hox gene tree analysis, we augmented a multiple sequence alignment based on our recent investigation of the embryonic transcriptome of a whip spider (Amblypygi, another member of Pedipalpi; Gainett and Sharma, 2020), by adding sequences of *M. giganteus*, the harvestman *Odiellus spinosus*, and the onychophoran *Euperipatoides kanangrensis*. For Nk, Iroquois/HRO, and SINE genes, we retrieved homologs for *I. scapularis* (tick) and *Centruroides sculpturatus* (scorpion) sequences from a recent comparative analysis of chelicerate homeobox genes (Aase-Remedios et al., 2023). Multiple sequence alignment was performed using Clustal-Omega (Sievers et al., 2011), followed by trimming of both ends to correspond to the previously identified conserved blocks (Gainett and Sharma, 2020; Aase-Remedios et al., 2023). Gene trees were inferred with a maximum likelihood analysis with IQ-TREE v.2 under a LG+I+G substitution model and 10,000 ultrafast bootstrap replicates (Le et al., 2012; Minh et al., 2013; Nguyen et al., 2014). After confirmation of orthology, gene locations were visually examined using gff files in Geneious Pro. Given extensive and recent explorations of these gene clusters for Arachnopulmonata (Gainett and Sharma, 2020; Harper et al., 2021; Ontano et al., 2021; Aase-Remedios et al., 2023; Sharma, 2023a), we mapped the relative locations of all identified genes with respect to subsets of selected species. We thus compared the architecture of *M. giganteus* Hox clusters to the chromosomal-level assemblies of *P. tepidariorum*, *I. scapularis*, and *O. spinosus*. We compared the architecture of the remaining clusters to *P. tepidariorum* and *I. scapularis*.

### 2.6 Synteny

For comparative analyses of macrosynteny, genomic positions of orthologs belonging to Bilaterian-Cnidarian-Sponge (BCnS) ancestral linkage groups (ALGs) defined by Simakov et al. (2022) were identified in the assembly of *Mastigoproctus giganteus* and visualized in Oxford dot plots using odp v.0.3.0 (Schultz et al., 2023). Orthologs were inferred between species by finding reciprocal-best protein matches using diamond (Buchfink et al., 2015) and OrthoFinder v.2.3.7 (Emms and Kelly, 2019). The reciprocal-best BLASTp hits were used to identify macrosyntenic chromosomes between species by performing Bonferroni-corrected one-sided Fisher’s exact tests (Simakov et al., 2020). We used in-built tools in odp v.0.3.0 to visualize syntenic patterns as Oxford dot plots and ribbon plots.

For analyses of self-synteny, collinearity within the *M. giganteus* genome was performed using One Step MCScanX feature of TBtools-II v.2.136 (Chen et al., 2023). Our search parameters consisted of BLAST hits = 5, evalue = ^1-10^, MATCH_SCORE = 50, MATCH_SIZE = 5, GAP_PENALTY = -1, OVERLAP_WINDOW = 5, and MAX GAPS = 25. Collinear blocks from the genome guided by the gff structural annotations from BRAKER3 were visualized using the advanced circos function in TBtools-II.

## Results and Discussion

### 3.1 Genome assembly and annotation

PacBio sequencing on two SMRTbells generated 182,391,985 and 176,172,263 high-quality long reads, totaling 98,345,435,739 bp, with read length N50 of 10,884 and 10,848 Kbp. Initial assembly of these reads with *hifiasm* resulted 3,825 contigs spanning 3,352,310,315 bp (contig N50: 3,581,478 bp), or 29.3× coverage. The purging of haplotigs from this assembly recovered 1,666 contigs spanning 3,166,202,185 bp (contig N50: 3,778,698 bp). Illumina sequencing of the Omni-C library generated 775 M 2×150 bp read pairs totaling 234 Gb, constituting 73.9× coverage of the PacBio haploid assembly. After scaffolding, the haploid assembly for *M. giganteus* was comprised of 3.17 Gb (scaffold N50: 236 Mbp; scaffold N90: 176 Mbp), with 31.1× coverage of PacBio CCS reads. The final assembly consisted of 185 scaffolds ranging in length from 12.4 Mbp to 28.1 Mbp (Fig. 2A). Fourteen scaffolds presumed to correspond to chromosomes represented 99.08% of the assembly length; the remaining 171 scaffolds represented 0.92% of the assembly length (Fig. 2B).

**Figure 2.**
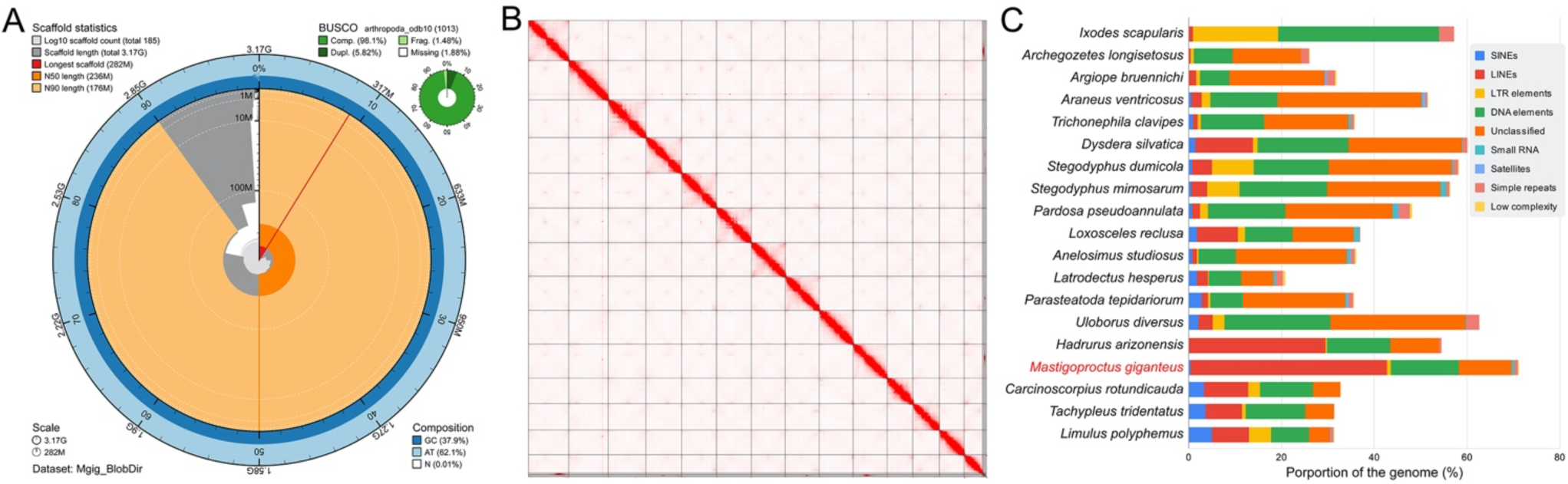
(A) Snail plot of *M. giganteus* scaffolded haploid genome assembly. (B) Hi-C contact map for the haploid assembly. Approximately 99% of data are scaffolded onto 14 pseudochromosomes. (C) Composition of chelicerate genome assemblies.

Analysis of genome completeness with the arthropod BUSCO dataset revealed the genome to be 98.1% complete; 5.82% of arthropod BUSCO genes were duplicated and 3.36% were fragmented or missing (Fig. 2A). Prior to structural annotation, RepeatMasker softmasked 2.26 Gbp (71.34%) of the genome. Repetitive elements occupy about 71.34 % of the *M. giganteus* genome with interspersed repeats comprising 69.8% of these regions. LINEs and SINEs make up 42.4% and 0.37% of the genome content whereas 11.4% of it are unclassified repeats. DNA transposons make up 14.6% of the genome (Fig. 2C). These values add to the substantial variation in genome composition previously reported for chelicerates (Grbić et al., 2011; Sanggaard et al., 2014; Hoy et al., 2016; Schwager et al., 2017; Shingate et al., 2020a; Gainett et al., 2021; De et al., 2023; Nuss et al., 2023). The high proportion of LINEs in the *M. giganteus* genome with respect to the remaining genomic elements mostly closely resemble the genome composition of the scorpion *H. arizonensis* (Fig. 2C).

We annotated the assembly with 95.8 Gbp of RNAseq read data generated from the same specimen, which exhibited a mean quality Phred score of 36.1 and mean GC content of 40.5%. Annotation of the genome with BRAKER yielded 47,271 transcripts corresponding to 41,197 putative gene models. Alignment with HISAT2 yielded a mapping rate of 87.04%. To improve precision of these gene models, we performed a PFAM filtration using the *hmmscan* function in HMMER v.3.4 (www.hmmer.org) against the PFAM database, retaining only gene models that retrieved hits. The final gene set consisted of 16,043 genes, which accords with known gene complements across Chelicerata.

### 3.2 Ohnologous gene clusters are consistently retained in the *M. giganteus* genome

Recent analysis of chromosomal-level genomes for spiders have shown that spiders broadly retain two Hox clusters, with each cluster found on a different chromosome (Fan et al., 2021; Aase-Remedios et al., 2023; Miller et al., 2023). Here, we expanded upon this analysis to examine the Hox cluster architecture of *M. giganteus* and the recently released chromosomal level assembly of the scorpion *Hadrurus arizonensis* (Bryant, 2024). Consistent with the inference of a shared duplication event, we found 15 Hox genes organized into two clusters, with each cluster on different pseudochromosomes (Fig. 3A). The first cluster (scaffold 1) spanned 2.03 Mbp and retained homologs of all Hox genes except *Hox3* and *Deformed*. The second cluster (scaffold 14) spanned 1.67 Mbp and retained homologs of all Hox genes except *Hox3*, *Ultrabithorax*, and *AbdominalB*. By comparison, apulmonate arachnids (*I. scapularis*, *O. spinosus*), retained a single Hox cluster, albeit with a partial and unannotated fragment of *abdominalA* in the assembly of the harvestman *O. spinosus*. Losses of duplicated Hox genes are common in the wake of whole genome duplication, as exemplified by the fragmentary clusters of horseshoe crabs (Shingate et al., 2020a, 2020b). Nevertheless, we suspect that the absence of some Hox genes in the *M. giganteus* and *O. spinosus* assemblies likely stem from insufficient mRNA sequencing data for annotation, given that orthologs of these missing genes are present in the developmental transcriptomes of Pedipalpi and Opiliones species that are better studied for developmental genetics (Sharma et al., 2012; Gainett and Sharma, 2020; Gainett et al., 2021). Indeed, duplicates of arachnopulmonate Hox genes tend to be strongly transcriptionally active during embryogenesis and are reliably amplified from cDNA of embryonic stages (Sharma et al., 2014b; Schwager et al., 2017), but we were unable to sample embryonic tissue from *M. giganteus* during the span of this study. Despite the gaps in the present assembly and the absence of embryonic transcriptome data, the retention of two Hox clusters on separate chromosomes supports the utility of this dataset for identifying shared genomic conditions across arachnopulmonates.

**Figure 3.**
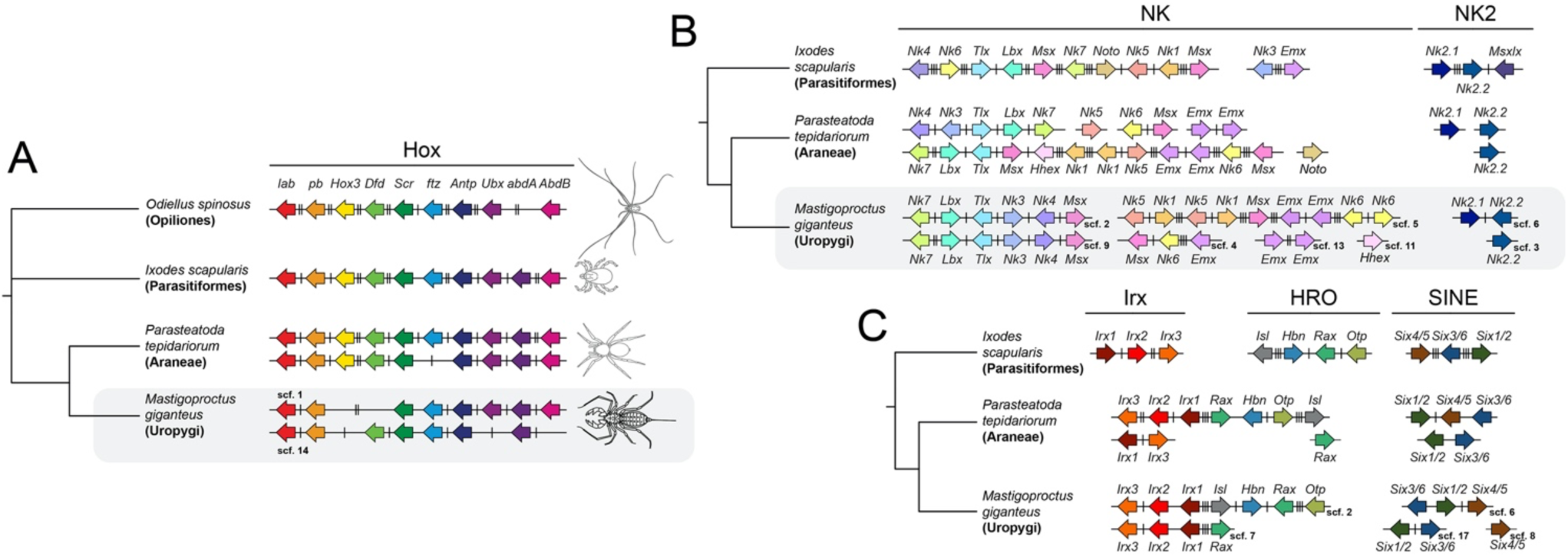
(A) Two ohnologous Hox clusters occur in *M. giganteus*. Colors correspond to individual Hox genes. (B) Two ohnologous Nk clusters occur in *M. giganteus*; the Nk2 cluster is disintegrated. (C) Two ohnologous Irx, HRO, and SINE clusters occur in *M. giganteus*. Note the frequent disintegration of one cluster in every pair. Distances between genes are indicated by hash marks: no hash marks indicate a distance less than 100 Kbp; one hash mark indicates a distance between 100 Kbp and 500 Kbp; two hash marks indicate a distance between 500 Kbp and 1 Mbp; and three hash marks indicate a distance greater than 1 Mbp. Figure conventions are based on Aase-Remedios et al. (2023) to enhance comparability.

Homologs of the Nk gene family exhibited more dynamic evolution, occurring on eight pseudochromosomes (Fig. 3B). Notably, we identified a block of the same five genes (*Nk7*, *Lbx*, *Tlx*, *Nk3*, *Nk4*, and *Msx*) occupying two pseudochromosomes, scaffold 2 (spanning 1.48 Mbp) and scaffold 9 (spanning 2.75 Mbp), with a single change in polarity in the *Nk4* homolog of the latter cluster. The same arrangement of five genes occurs on separate chromosomes of multiple spider species. A less organized cluster occurring on scaffold 5 (spanning 63.6 Mbp) contained tandem duplicates of *Nk1*, *Nk5*, *Nk6*, *Emx*, and a single homolog of *Msx*. This closely parallels the retention of the same set of tandemly duplicated genes in spiders (Aase-Remedios et al., 2023), with the exception that the *Nk6* tandem duplication is restricted to *M. giganteus*. The *Nk6* tandem duplication may be specific to Pedipalpi or the second copy of *Nk6* in this cluster may have been lost in the common ancestor of spiders. We also found a second linkage group of *Msx* and *Nk6* in scaffold 4 and two copies of *Emx* in scaffold 13, both of which closely reflect the condition found in spiders. These shared patterns suggest that the *Msx-Nk6* duet predated the divergence of Tetrapulmonata. The Nk2.2 cluster was closely comparable to those reported in spider genomes as well. We did not find a homolog of *Msxlx* in the *M. giganteus* genome; its widespread absence across spiders suggests two independent losses of *Msxlx* in Pedipalpi and in the common ancestor of entelegyne spiders.

Iroquois genes have a complex evolutionary history. It is inferred that three tandem duplicates were formed in the common ancestor of Arthropoda from a single *Irx* ancestral gene, with subsequent loss of *Irx3* in the common ancestor of Pancrustacea (Kerner et al., 2009; Setton et al., 2024). *Irx1* underwent another tandem duplication in a subset of insects to form the well-studied genes *araucan* and *caupolican*, which play key roles in insect embryogenesis, together with *Irx2* (*mirror*) (del Corral et al., 1999; Cavodeassi et al., 2001). Due to whole genome duplication, up to six copies of *Irx* genes occur in spiders, with one of these (*Irx3-2* or *waist-less*) recently shown to act as a gap segmentation gene in *P. tepidariorum* (Setton et al., 2024). In *M. giganteus*, we found two clusters comprised of the three Iroquois homologs, with identical arrangement of gene polarity and similar linkage to genes in the HRO cluster (Fig. 3C). One pseudochromosome (scaffold 2) contained all three *Irx* genes as well as the LIM-class gene *Islet* and the PRD-class genes *Homeobrain*, *Retinal homeobox* (*Rax*), and *orthopedia* (*otp*). The second cluster (scaffold 7) contained the three *Irx* genes and the second copy of *Rax*. The organization of the first and larger cluster closely parallels the arrangement of these genes in the unduplicated genome of *I. scapularis*, as well as the larger “A” cluster of many spiders wherein Irx and HRO are linked. This pattern of occurrence suggests that an Irx-HRO linkage reflects the ancestral organization of Tetrapulmonata. By contrast, the spider “B” cluster is more dynamic, with the second *Rax* homolog often not found on the same chromosome as the second *Irx* cluster (Aase-Remedios et al., 2023).

The architecture of the *M. giganteus* SINE clusters consists of a single cluster containing *Six1/2*, *Six3/6*, and *Six4/5* in proximity on one pseudochromosome (scaffold 6, spanning 0.33 Mbp), a second cluster containing *Six1/2* and *Six3/6* on another pseudochromosome (scaffold 17, spanning 0.22 Mbp), and the second paralog of *Six4/5* on a third scaffold (scaffold 8) (Fig. 3C). The retention of two copies of each of the three *Six* homologs present in the arthropod common ancestor is consistent with the inference of whole genome duplication. In addition, the presence of two *Six4/5* copies in *M. giganteus* supports the previous inference that the loss of the second *Six4/5* homolog occurred after the divergence of Araneae from the remaining tetrapulmonates (Aase-Remedios et al., 2023).

Taken together, the incidence of these surveyed genes as pairs of clusters on different chromosomes in *M. giganteus* is broadly consistent with the inference of a shared whole genome duplication in the common ancestor of arachnopulmonates. Notably, the fidelity of gene order after duplication varies dramatically from one cluster to the next, with the greatest conservation in the well-studied Hox genes and the least in the Nk cluster.

### 3.3 Macrosynteny analyses reveal the signature of whole genome duplication

Despite accumulating evidence of conserved gene clusters in the genomes of various arachnopulmonates, a recent work has questioned the validity of previous evidence for the arachnopulmonate whole genome duplication, concluding that there was insufficient evidence for this event. Thomas et al. (2024) contended that much of the support for the arachnopulmonate WGD relies only on a small portion of the genome, with emphasis on the shared presence of duplicated Hox clusters. While it is true that early reports of this duplication event emphasized the Hox cluster, which is known to serve as a reliable readout of Paleozoic genome duplications (Wagner et al., 2003; Dehal and Boore, 2005; Crow et al., 2006; Schwager et al., 2007, 2017; Sharma et al., 2014b; Shingate et al., 2020b), subsequent analyses have since documented widespread retention of paralogs within many key developmental pathways in arachnopulmonates, to the exclusion of apulmonate taxa. Gene families and pathways exhibiting systemic arachnopulmonate duplications include the entire homeobox family (Leite et al., 2018; Ontano et al., 2021; Aase-Remedios et al., 2023; Gainett et al., 2023), proximodistal axis patterning genes (Gainett and Sharma, 2020; Nolan et al., 2020; Ontano et al., 2021; Klementz et al., 2024), microRNAs (Leite et al., 2016; Ontano et al., 2021), segmentation and posterior axis patterning genes (Benton et al., 2016; Bonatto Paese et al., 2018; Baudouin-Gonzalez et al., 2021; Harper et al., 2021; Janssen et al., 2021; Setton and Sharma, 2021), and the retinal determination network (Samadi et al., 2015; Schomburg et al., 2015; Gainett et al., 2020, 2024).

The systemic duplication and retention of these genes across arachnopulmonate orders, together with highly suggestive gene tree topologies (Nolan et al., 2020; Ontano et al., 2021), render it highly unlikely that these patterns of paralogy occurred via multiple and independent duplications. We aimed to understand how different studies could reach opposite conclusions regarding the arachnopulmonate duplication. Thomas et al. (2024) undertook three analytical approaches to testing for a shared genome duplication event: gene tree reconciliation, analysis of distributions of synonymous substitutions per site between paralogs (K_s_ distances), and tests for self-synteny. They found no evidence to support a WGD in the common ancestor of Arachnopulmonata using these tests; they found evidence of only a single lineage-specific duplication within Xiphosura (horseshoe crabs), despite previous inferences of three rounds of duplication in this group (Kenny et al., 2016; Shingate et al., 2020a; Nong et al., 2021).

There are, however, important caveats in the use of the methods implemented by Thomas et al. (2024) for analysis of ancient genome duplications. Gene tree reconciliation methods rely on embedding a gene tree into an underlying species tree, minimizing mismatches via optimization of gene duplication and loss events. Crucially, these methods require a well-resolved gene tree topology, with lower bootstrap support values at particular nodes leading to more extreme bias (Hahn, 2007). However, gene tree topologies with high support values can still erroneously represent the evolution of orthologs, particularly in cases of gene duplication followed by functional divergence. In cases of gene neofunctionalization, one gene copy is selectively constrained by retention of an ancestral function (with an ensuing lower rate of evolution), whereas its sister copy acquires a new function and exhibits a higher rate of evolution (particularly in cases where a new functional domain is acquired; Holland et al., 2017). The more rapid rate of evolution of the second copy can yield artefactual clustering, due to a *de facto* long-branch attraction artifact at the level of the gene tree (Swenson and El-Mabrouk, 2012). Moreover, the pseudogenization and loss of numerous duplicated genes is anticipated to the be most likely outcome of ancient whole genome duplication events, as exemplified by the two rounds of genome duplication that occurred at the base of the vertebrate tree of life (Dehal and Boore, 2005; Simakov et al., 2020, 2022). Likewise, the use of K_s_ plots to visualize the distribution of synonymous divergence between duplicated paralogs is known to be inadequate for detection of ancient (i.e., pre-Cenozoic) whole genome duplication events, given routine failure when less than 10% of paralogs are retained (Tiley et al., 2018). K_s_ plots also exhibit sensitivity to gene saturation, which may violate the assumptions of a simple nucleotide substitution model over long spans of evolutionary time (Rabier et al., 2014). These considerations bear heavily upon the analysis of the arachnopulmonate whole genome duplication, which is at least Silurian in age, with *bona fide* stem-group fossils of scorpions (and therefore, crown-group arachnopulmonates) present by 430 Myr (Waddington et al., 2015). Molecular clock estimates for Chelicerata have implied that the radiation of arachnopulmonates may represent the oldest known whole genome duplication in the animal tree of life (Lozano-Fernandez et al., 2016; Ballesteros et al., 2021). Traditional tests of whole genome duplication (e.g, gene tree reconciliation and K_s_ plots) may not be able to accommodate the noise incurred by such ancient divergences, despite their successful application in cases of phylogenetically recent whole genome duplication events (e.g., angiosperms; snails; Tang et al., 2010; Simakov et al., 2020; Farhat et al., 2023).

It is nevertheless surprising that Thomas et al. (2024) found little or no evidence for self-synteny in arachnid genomes, given that recent efforts had already previously reported patterns of self-synteny in spider species (Fan et al., 2021; Miller et al., 2023). We suspect that this discrepancy could be explained by the stringent definition of synteny employed by Thomas et al. (2024); they defined a syntenic block as a group of at least three genes present in the same order in multiple regions of the genome. As shown by our analysis of the Nk cluster (Fig. 3B), the emphasis on conservation of gene order can mask true patterns of gene linkage when even minor rearrangements occur within a syntenic block. However, a recently developed suite of methods for detecting ancient syntenic patterns embraces instead the classical definition of synteny: gene linkage without regard to a specific gene order. This approach has unveiled widespread signatures of ancient bilaterian chromosomes (Simakov et al., 2020, 2022; Schultz et al., 2023) and proven especially successful in detecting multiple rounds of WGD in the evolutionary history of the vertebrates (Schultz et al., 2023).

We therefore analyzed selected chelicerate genomes using the odp v.0.3.0 suite, with the BCnS (Bilateria-Cnidaria-and-Sponge) ancestral linkage groups (ALGs), which consists of 29 ancestral chromosomal blocks inferred to have been present in the common ancestor of Myriazoa. The genome of the tick *I. scapularis* and the mite *Sarcoptes scabiei* revealed no signature of systemic genome duplication. (Fig. 4A, 4B). In the tick (Fig. 4A), significant numbers (> 5 genes; *p* < 0.05) of 28 ALGs were associated with single chromosomes; only ALG Ea was found with significant numbers of genes on two chromosomes. In the mite (Fig. 4B), a larger proportion of non-significant hits was found throughout the genome, reflecting a higher rate of genome evolution previously reported for many miniaturized, parasitic Acariformes and Parasitiformes (Hoy et al., 2016; Pace et al., 2016). Nevertheless, 27 ALGs were associated with single chromosomes; only ALG Ea and H exhibited significant numbers of hits on two chromosomes.

**Figure 4.**
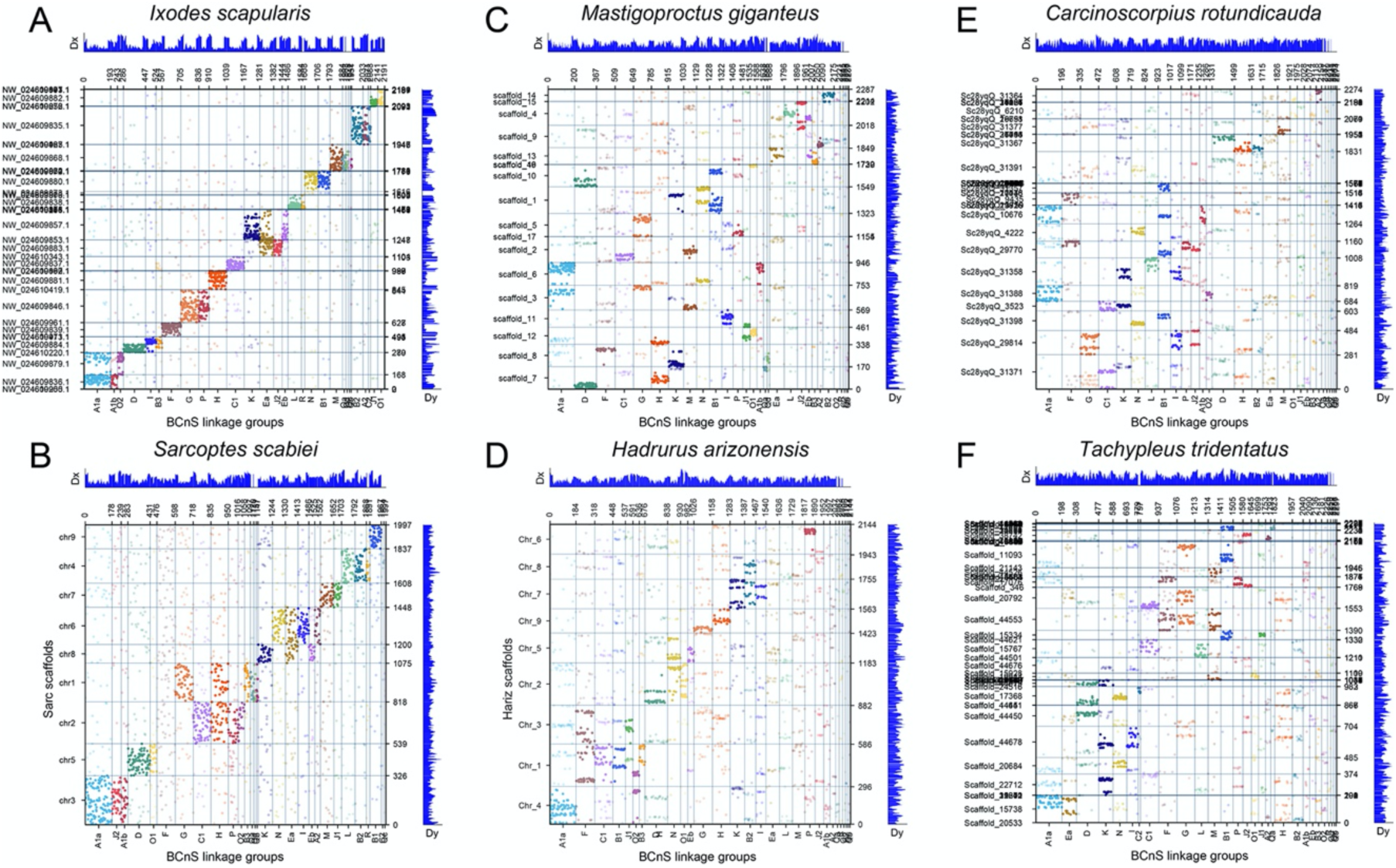
Oxford dot plots showing the locations of BCnS homologs in selected chelicerate genomes. Colors correspond to individual ancestral linkage groups. Solid circles indicate significant hits (*p* < 0.05); transparent circles indicate insignificant hits. (A) The tick *Ixodes scapularis*, unduplicated genome. (B) The mite *Sarcoptes scabiei*, unduplicated genome. (C) The vinegaroon *Mastigoproctus giganteus*, 1R whole genome duplication. (D) The scorpion *Hadrurus arizonensis*, 1R whole genome duplication. (E) The horseshoe crab *Carcinoscorpius rotundicauda*, 3R whole genome duplication. (F) The horseshoe crab *Tachypleus tridentatus*, 3R whole genome duplication.

We compared these patterns to two arachnopulmonates (*M. giganteus* and the scorpion *Hadrurus arizonensis*), as well as two horseshoe crabs (*C. rotundicauda* and *T. tridentatus*). In *M. giganteus* (Fig. 4C), 12 of the 29 ALGs were represented on two separate chromosomes, 13 ALGs had significant hits to a single chromosome (albeit with frequent signatures of non-significant hits in other chromosomes), and four ALGs exhibited only non-significant hits. The genome of the scorpion (Fig. 4D) exhibited a noisier pattern, with four ALGs exhibiting significant hits to two chromosomes, 13 ALGs associated with a single chromosome, and 12 with no significant hits. As with *M. giganteus*, numerous non-significant hits were found for the ALGs occurring on single chromosomes, often in numbers suggestive of systemic duplication. Notably, only one ALG exhibited shared duplication in *M. giganteus* and *H. arizonensis* (ALG N), which may reflect lineage-specific patterns of gene retention since the divergence of these taxa in the Silurian (or earlier).

In the horseshoe crabs, at least one of the whole genome duplications is thought to be recent, as it exhibits a clear signature of paralogy in K_s_ plots (Roelofs et al., 2020). In *C. rotundicauda*, we found one ALG with genes on four scaffolds, seven ALGs with genes on two scaffolds, 10 ALGs present on only one chromosome, and 11 ALGs with no significant mapping (Fig. 4E). In *T. tridentatus*, we found five ALG with genes on three scaffolds, four ALGs with genes on two scaffolds, 10 ALGs present on only one chromosome, and 10 ALGs with no significant mapping (Fig. 4F). Six ALGs (B1, C1, F, J2, K, N) exhibited significant retention of genes on more than one chromosome for both species. As with the arachnopulmonates, this pattern may reflect lineage-specific retention of paralogs in the wake of the last whole genome duplication.

Notably, the signature of ancient genome duplication elicited through this investigation recapitulates the same dynamics previously found in vertebrate genomes. In comparisons of tetrapod (twofold genome duplication) and teleost fish (threefold genome duplication) genomes against a cephalochordate (unduplicated), (Simakov et al., 2020) found few cases of ALGs that faithfully mapped to four or eight chromosomes in the tetrapod and telost, respectively. This trend reflects the phenomenon of common and rapid gene losses in the wake of genome duplications. For example, only 5% of genes in the human genome comply with the 4:1 rule, i.e., the retention of exactly four ohnologs in human to one single-copy homolog in an invertebrate outgroup (Friedman and Hughes, 2003; Dehal and Boore, 2005; Inoue et al., 2015).

Given the relative cohesion of the *I. scapularis* genome with respect to the ancestral myriazoan linkage groups, we visualized comparisons of selected genomes against this benchmark (Fig. 5). Despite the evolutionary distance between these taxa, we found several genomic regions that harbored shared ALGs for *I. scapularis* and *S. scabiei* (Fig. 5A), without evidence of systemic duplication. By contrast, comparison of the *M. giganteus* and the *T. tridentatus* genomes against the *I. scapularis* reference showed several syntenic blocks that were on a single chromosome in the tick, but present in two chromosomes in the vinegaroon (Fig. 5B) and in two to four chromosomes of the horseshoe crab (Fig. 5C). This is readily visualized via ribbon plot, with mapping of *I. scapularis* genes to their paralogs in the *M. giganteus* genome (Fig. 5D). This pattern of localization of duplicated genes on different chromosomes echoes visulizations of duplicated genes in spiders (Aase-Remedios et al., 2023) and in horseshoe crabs (Shingate et al., 2020a). Lastly, contrary to the previous analysis that did not recover self-synteny in arachnid genomes, we discovered 30 syntenic blocks with duplicates on other chromosomes in *M. giganteus* and 13 blocks with duplicates on the same chromosome. The discovery of self-syntenic patterns in this arachnopulmote reflecting patterns previously reported for spiders (Fan et al., 2021; Miller et al., 2023) (Fig. 5E).

**Figure 5.**
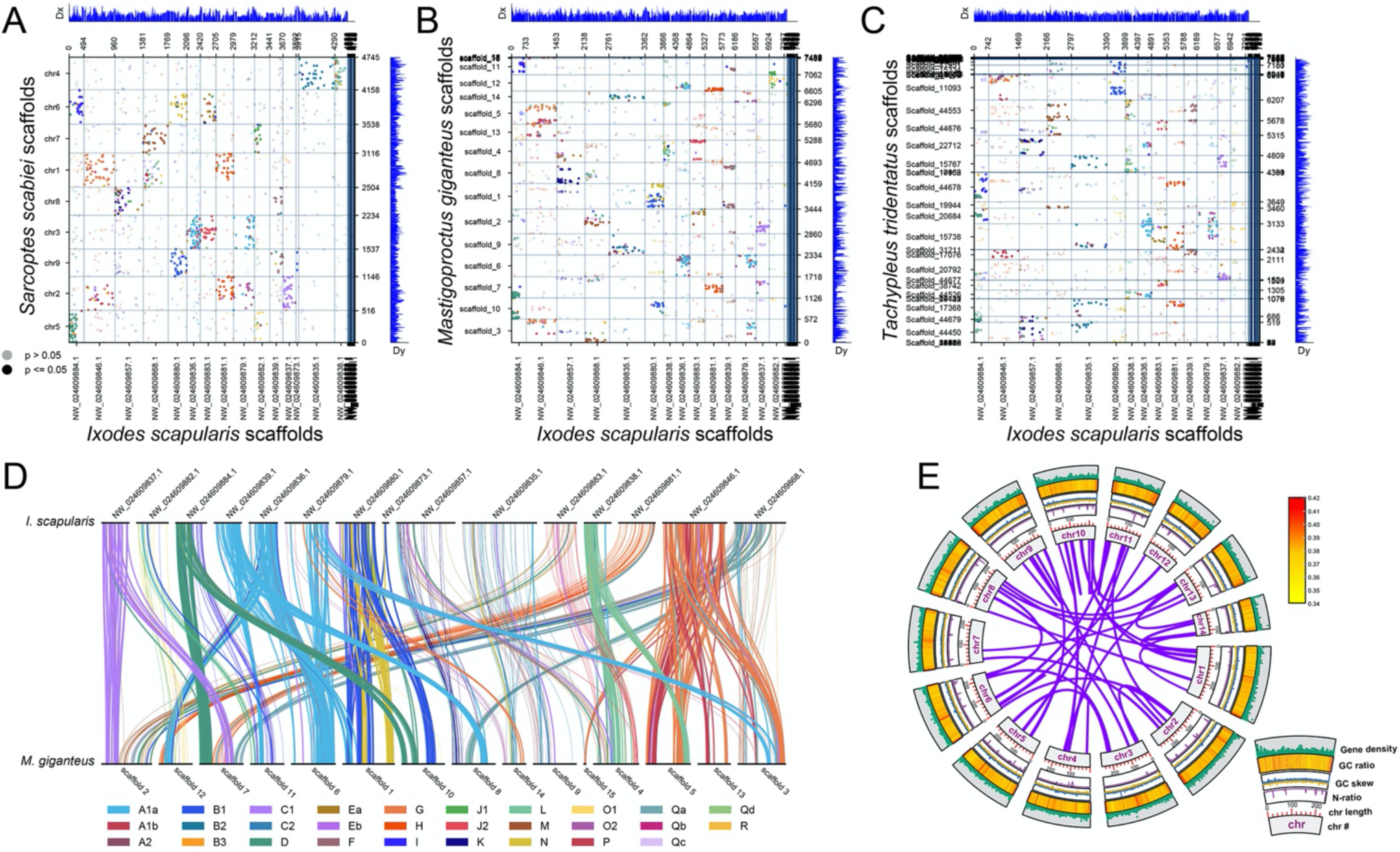
(A-C) Oxford dot plots of species-by-species comparisons, based on the BCnS gene set. Colors correspond to individual ancestral linkage groups. Solid circles indicate significant hits (*p* < 0.05); transparent circles indicate insignificant hits. (A) *Sarcoptes scabiei* against *Ixodes scapularis*. (B) *Mastigoproctus giganteus* against *Ixodes scapularis*. (C) *Carcinoscorpius rotundicauda* against *Ixodes scapularis*. (D) Ribbon plot of BCnS homologs in *I. scapularis* and *M. giganteus*. Each line corresponds to an orthologous pair. Lines are thickened for genes in ancestral linkage groups that exhibit duplications, to visualize their appearance on two separate chromosomes in *M. giganteus*. (E) Self-synteny in the *M. giganteus* assembly. Each band represents a syntenic block found in the same configuration in another genomic location.

Taken together, these results suggest that the signature of an ancient and shared whole genome duplication that unites Arachnopulmonata is systemic. This single event most parsimoniously explains the pervasive paralogy of genes, miRNA families, gene tree topologies, syntenic blocks, and shared expression patterns of daughter copies independently validated by an array of workers.

## 4 Conclusion

The availability of a chromosomal-level assembly for a member of the clade Pedipalpi unlocks the phylogenetic polarization of genomic events within Arachnopulmonata, with emphasis on understanding genomic dynamics in spiders. The conserved organization of the vinegaroon genome, as compared to spider and scorpion counterparts, offers numerous advantages for reconstructing ancestral genomic conditions within this group of arachnids.

## Author Contributions

SSK: Conceptualization, PacBio and HiC library preparation and sequencing, genome assembly, genome annotation, bioinformatic analysis of synteny. BCK: Taxonomic identification, specimen preparation, RNA sequencing, bioinformatic analysis of gene clusters, writing. PPS: Conceptualization, funding acquisition, bioinformatic analysis of gene clusters, writing.

## Acknowledgements

PacBio sequencing was performed at the University of California-Irvine Genomics Research and Technology Hub. Illumina sequencing for transcriptomes and Omni-C libraries was performed at the University of Wisconsin-Madison Biotechnology Center. Kaitlyn M. Abshire, Hugh G. Steiner, Sophie M. Neu, and Ethan M. Laumer assisted with the care and maintenance of *M. giganteus*. Bioinformatic analyses were conducted on the Colonial One High Performance Computing Facility at The George Washington University facilitated by Gustavo Hormiga.

## Data availability

The final genome assembly, raw data from the PacBio library, raw data from the Hi-C library, raw data from RNAseq, and the annotation file have been deposited at NCBI as a single BioProject. Supplementary materials consisting of the final gene set, final peptide set, multiple sequence alignments, gene tree topologies, and outputs of odp v.0.3.0 have been deposited in FigShare.

